# Multi-Funnel Landscape of the Fold-Switching Protein RfaH-CTD

**DOI:** 10.1101/221143

**Authors:** Nathan A. Bernhardt, Ulrich H.E. Hansmann

## Abstract

Proteins such as the transcription factor RfaH can change biological function by switching between distinct three-dimensional folds. RfaH regulates transcription if the C-terminal domain folds into a double helix bundle, and promotes translation when this domain assumes a *β*-barrel form. This fold-switch has been also observed for the isolated domain, dubbed by us RfaH-CTD, and is studied here with a variant of the RET approach recently introduced by us. We use the enhanced sampling properties of this technique to map the free energy landscape of RfaH-CTD and to propose a mechanism for the conversion process.

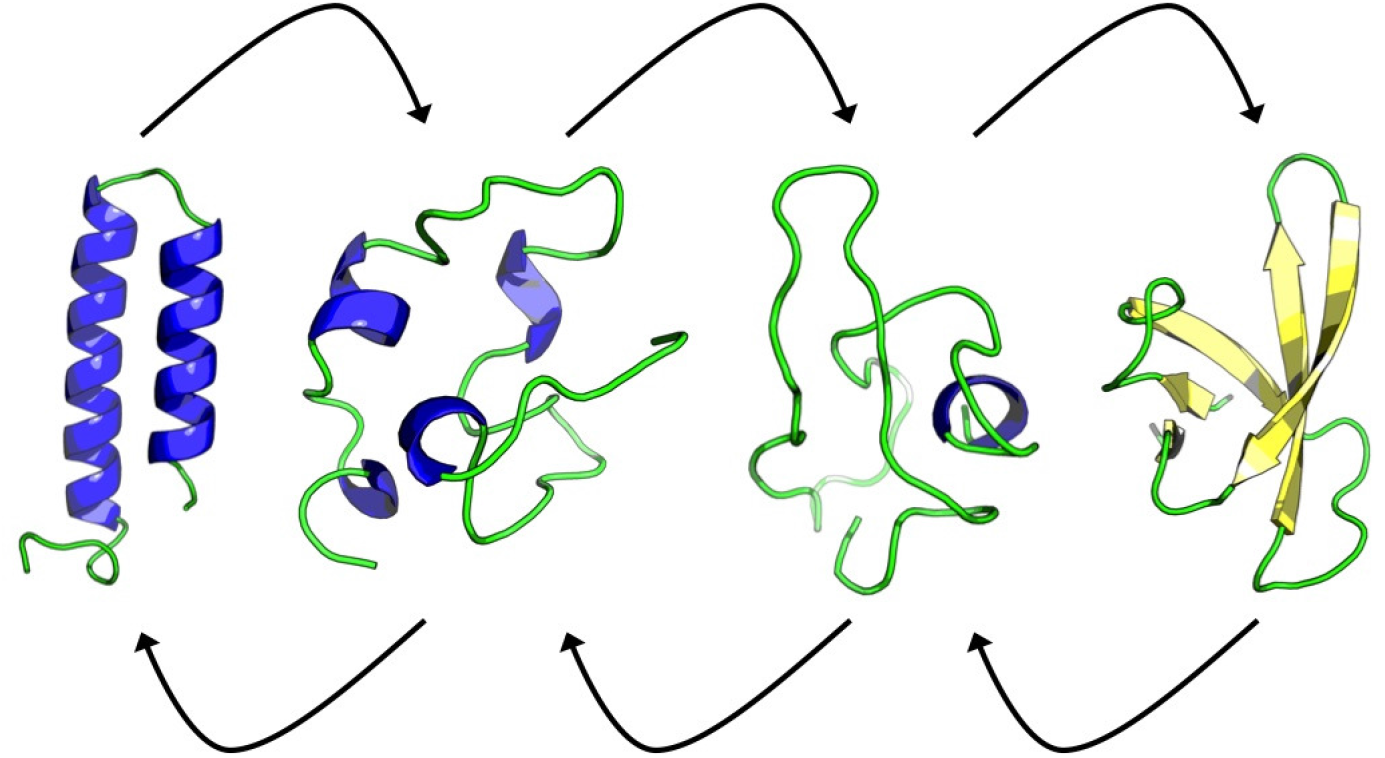
TOC Image

## Introduction

Proteins play a central role in the biochemistry of cells, participating in transcription, cell signaling, migration and muscle movement to name only a few of their roles. Protein function is correlated with the molecule assuming a specific three-dimensional shape, but the process by that a protein folds into a certain structure is not known in all details, and depends not only on the sequence of amino acids (the chemical composition of a protein) but also on environment and interaction with other molecules. In the standard model of protein folding, one assumes that a protein has a funnel-shaped energy landscape^1,2^ that guides a multitude of possible folding pathways into a unique structure where the protein is biologically active. However, such a single-funnel picture cannot describe all aspects of folding for all proteins. For instance, intrinsically disordered proteins^3-5^ do not have a defined structure, but may assume one when interacting with other proteins, with the structure changing with the binding partner. In other cases, proteins exist in an ensemble of different (but defined) structures^6-8^ which may allow proteins to have more than one function in the cell. In these cases we would expect a multi-funnel shaped folding landscape.

Take as an example RfaH,^9^ a protein that triggers gene expression in Escherichia coli by switching the structure of the C-terminal domain from an *α*-helical hairpin (PDB-ID: 2OUG) to a *β*-barrel (PDB-ID: 2LCL), see Figure 1. In the first form, stabilized by interaction with the N-terminus, the C-terminal domain masks a RNA polymerase binding site on the N-terminal domain thus regulating transcription. However, when not in contact with the N-terminal domain, or as an isolated protein, the C-terminal domain spontaneously rearranges into a *β*-barrel (figure 1). In this form, RfaH binds directly to the ribosomal protein S10, thus recruiting the prokaryotic ribosomal 30S subunit to the elongating RNA and promoting translation. Hence, the fold switch of the C-terminal domain alters dramatically the function of the RfaH protein. Both folds are encoded in the sequence of the C-terminal domain, and it is the interaction (or lack of interaction) with the N-terminus of RfaH that selects the fold. Hence, we would expect a double-funneled landscape for the isolated 66-residue large C-terminal domain of RfaH (RfaH-CDT), with one funnel leading to the *β*-barrel, and the secondary funnel leading to the *α*-helical hairpin.^10^ The rather small size, the experimentally observed fold switching, and the resolved structures of the two folds make RfaH-CTD an ideal model to study the factors that determine protein plasticity and the mechanism of fold switching in proteins.

**Figure 1:**
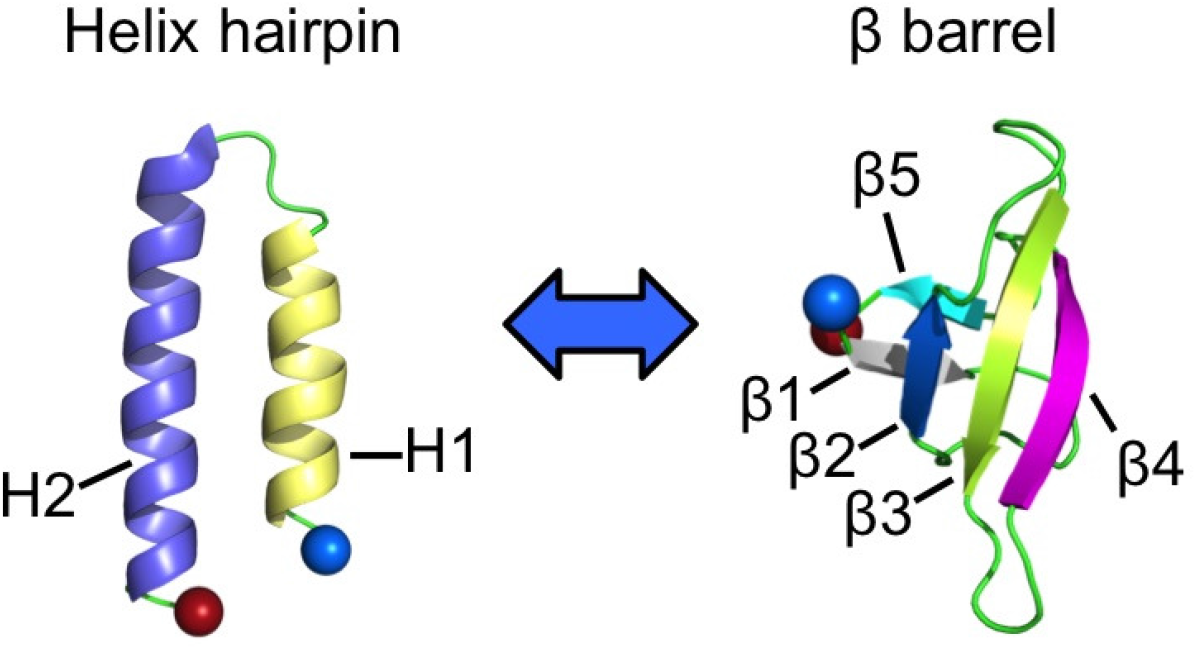
The two folds observed for the C-terminal domain (RfaH-CTD) of the transcription factor RfaH. The N-terminus of RfaH-CTD is marked by a blue ball and the C-terminus a red ball.

However, probing such fold switching and mapping their energy landscape by experiments or *in silico* is a challenge.^10-14^ Computationally, the problem is that the exploration of the ensemble of possible structures and the conversion between these structures happens on timescales that, on general-purpose computers, are not accessible in all-atom molecular dynamics simulations with explicit solvent. Enhanced sampling techniques such as Replica Exchange Molecular Dynamics (REMD)^15-20^ promise to overcome this problem by realizing a random walk in temperature which allows the system to escape out of traps and cross barriers by explorations to higher temperatures. However, the sampling efficiency of REMD is often below the theoretical maximum. One problem is that the probability for a replica exchange depends on the temperature spacing which shrinks dramatically with system size. Hence, with the inclusion of water molecules one needs even for small proteins a huge number of replica. This often makes REMD simulations of proteins with explicit solvent impractical, and previous REMD simulations of RfaH^11^ had for this reason to rely on an implicit solvent. While these simulations allowed the authors to propose a transition pathway between the two folds, choice of an implicit solvent is not without problems. While the helix-hairpin was found, the simulation was unable to completely fold the *β*-barrel. Only when continuing the simulations from the best configurations by including solvent molecules explicitly was the correct *β*-barrel fold found. ^11^

We have recently proposed to overcome some of the limitations that hold back REMD by a Replica-Exchange-with-Tunneling (RET) approach. ^21^ We have shown that RET in conjunction with a Hamilton-Replica-Exchange^22,23^ of systems where the “physical” system is coupled to varying degrees with biasing Go-models, allows efficient simulation of proteins and protein assemblies that can take more than one state.^24,25^ For instance, we have used this approach in a a recent study^24^ of the two mutants GA98 and GB98, which differ only in a single residues but keep the distinct original folds of the A and B domain of protein G, GA and GB.^26-28^ We have shown how the mutation leading from GA98 to GB98 alters the energy landscape leading to selection of one fold over the other. ^24^ In the present work we use the superior sampling properties of this approach to explore the folding and switching landscape of the C-terminal domain of RfaH, Rfah-CTD, and propose a transition pathway that connects the two forms.

## Materials and Methods

In its standard implementation, replica-exchange sampling^15,16,18^ aims to enforce a random walk in temperature as a way to escape out of local minima in order to achieve faster convergence at a (low) target temperature. However, the exchange move between neighboring temperatures often leads to a proposal state that is exponentially suppressed, but if accepted the multiple replica system quickly relaxes to a state of comparable probability to that before the exchange. As a way to overcome this bottleneck and tunnel through the unfavorable transition we have recently introduced Replica-Exchange-with-Tunneling (RET)^21,24,25^ by the following four-step-procedure:

1. In the first step, the configurations *A*(*B*) evolve on two neighboring replica over a short microcanonical molecular dynamics trajectory to configurations *A*′(*B*′), without that the total energies *E*_1_ and *E*_2_ change on the two replicas. However, there will be an exchange between potential and kinetic energy on each replica.
2. Next, the configurations *A*′ and *B*′ are exchanged, and the velocities are rescaled according to the following equations such that the energies remains constant before and after the exchange: 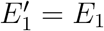 and 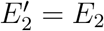.

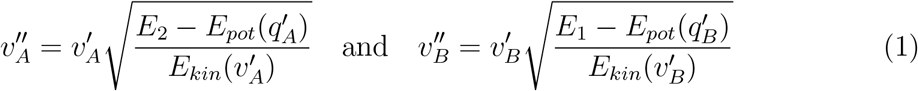
3. After the exchange, the two replica evolve again by microcanonical molecular dynamics. While the total energies *E*_1_ and *E*_2_ on the two replica do not change, the exchange between potential and kinetic energy will lead to final states 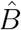 on replica 1 and *Â* on replica 2 that have potential energies comparable to the corresponding configurations before the exchange move, and velocity distributions as one would expect for the given temperatures at each replica.
4. The final configurations on each replica are now either accepted or according to the following Metropolis criterium

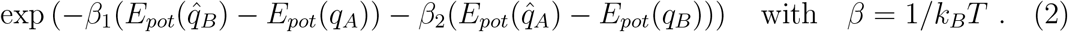 If rejected, molecular dynamics simulations continue with the original configurations *A*(*B*). However, in both cases, new velocity distributions are randomly drawn according to the temperatures on the respective replica.

For more details on this approach and its limitations, see Refs.^21,24^

When simulating conformational transitions in systems with competing attractors as in the case of RfaH-CDT (an *α*-helical hairpin and a *β*-barrel), one can combine the RET move with exchange moves between “physical” models and such relying on Go-type force fields that bias toward one or the other of the competing configurational states. Go-models have very smooth energy landscapes with a single funnel located around the target fold. As such, Go models fold proteins quickly, but are by construction unable to capture accurately the energetics of non-native folds. Thus, in an effort to exploit the quick folding of Go models but remove in our simulations the associated bias against non-native folds, one can think of a Hamilton Replica Exchange Method^22^ where at each replica a “physical” model is “fed” by a Go-model, but where the bias differs for each replica.^25,29,30^ In the present implementation, the physical and the Go-model are coupled through a potential that depends on the similarity between configurations in the two models^30,31^

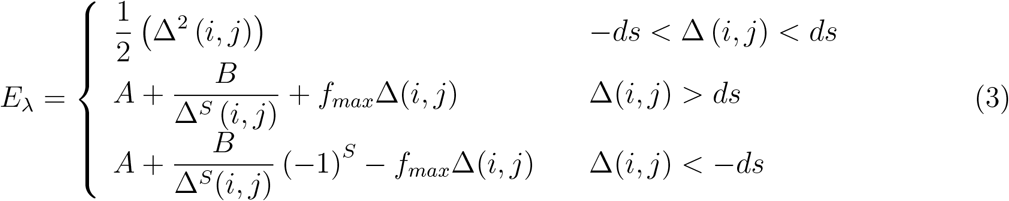

such that Δ (*i*, *j*) is the difference in distances between alpha carbons i and j in the respective models, *f_max_* is a parameter that controls the maximum force as Δ (*i*, *j*) → ∞, and S controls how fast this value is reached. The parameters A and B are set so that the potential and its first derivative are continuous at values of Δ (*i*, *j*) = ±*ds*, and are expressed as

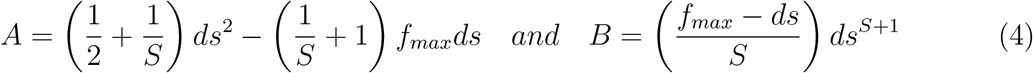

Thus the total potential energy of the system is

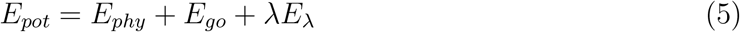

where λ controls how strongly the physical and Go-models are coupled. Hamilton Replica Exchange now introduces a random walk in λ-space, with data to be analyzed only from the replica where the bias from the Go-model on the physical model vanishes, i.e., λ = 0.

However, exchange rates are often low in such an approach, a problem that is avoided in our simulations by using RET moves, where now the velocities are rescaled according to

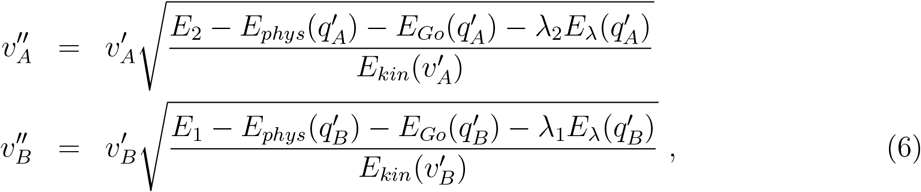

and RET moves are accepted with probability

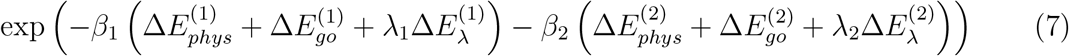

where 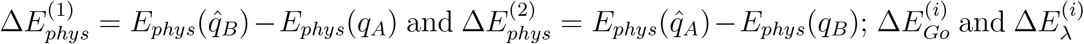 are defined accordingly.

In the present example we have two ladders of replica, each covering a range from λ = 0 to a value λ = λ_*max*_. In the one ladder, replicas walk between a system with no bias on the physical model to one where there is maximal bias toward the α-helix hairpin; in the other the biasing is toward the *β*-barrel. The two λ = 0 replica do not exchange configurations but serve as reservoirs from that a canonical simulation at the same temperature is “fed” by a heat bath move.

We consider in our simulations not the full-length RfaH protein but only the 48-residue-long C-terminal domain, RfaH-CTD. The RfaH-CTD protein is capped at the N-terminus by an acetyl group, and by a methylamine group at the C-terminus. Prior to running large scale simulations initial structures were randomized by high temperature molecular dynamics (T = 3500 K). Replicas were then brought to their initial lambda values in a short preproduction run. A total of 25 replicas with a lambda distribution of λ = 0.4, 0.2, 0.1, 0.075, 0.055, 0.035, 0.028, 0.023, 0.015, 0.010, 0.005, 0.00, 0.00, 0.00, 0.005, 0.010, 0.015, 0.023, 0.028, 0.035, 0.055, 0.075, 0.1, 0.2 and 0.4 is used. The *E*_λ_ energy function of Eq. 3 is parametrized with *ds* = 0.3Å, *S* = 1 and *f_max_* = 0. Data were generated over 100 ns trajectories using an in house version of GROMACS 4.6.5^32^ (available from the authors on request), modified to accommodate RET sampling and the Go-model feeding. Potential energy calculations relied on the CHARM36 force field in combination with a GBSA implicit solvent^33^ for the physical model and the smog energy function^34,35^ for the Go-model (using the online SMOG server at http://smog-server.org). Equations of motion are integrated by a leap frog integrator with a time step of 2 fs, which requires use of the linear constraint solver (LINCS)^36^ for constraining hydrogen and heavy atom bond distances. A plain cutoff of 1.5 nm was used for treatment of electrostatics, and the v-rescale thermostat^37^ is used to keep the temperature at 310 K.

## Results and Discussions

We start our analysis by first testing that our simulations have converged. For this purpose, we have calculated the later discussed free energy landscape for different intervals of the 100 ns trajectory, see Figure SF1 in the Supplemental Informations. Comparing these landscapes we see that our trajectory has converged after 30 ns, and therefore use the last 70 ns for our analysis. Within this time interval, we observe an average exchange rate between neighboring replicas, with individual rates listed in table ST1, of 26 ± 3%, a value that is similar to the one seen by us in previous RET simulations where we also showed that regular Hamilton Exchange Replica Exchange let to lower rates (especially around λ = 0) if the same number of replica is used. ^21,24,25^ As a consequence, replica can walk on both sides of the ladder between replica with λ = λ_*max*_ where the physical model is biased strongly by the corresponding Go-model, and λ = 0 where the physical model is not biased by the Go-model. The number of walks between the two extreme values (called by us tunneling events) are a measure for the quality of simulation. In the present study, we observe a total of 34 tunneling events, with examples shown in Figure 2, a value that in our previous work indicated that our simulations had sampled sufficient statistics. We remark that we saw in previous simulations^21,24,25^ always much higher numbers of tunneling events when using RET exchange moves than in regular Hamilton Exchange Replica Exchange with the same number of replicas and the same λ distribution, reflecting the superior sampling that results from the RET move.

**Figure 2:**
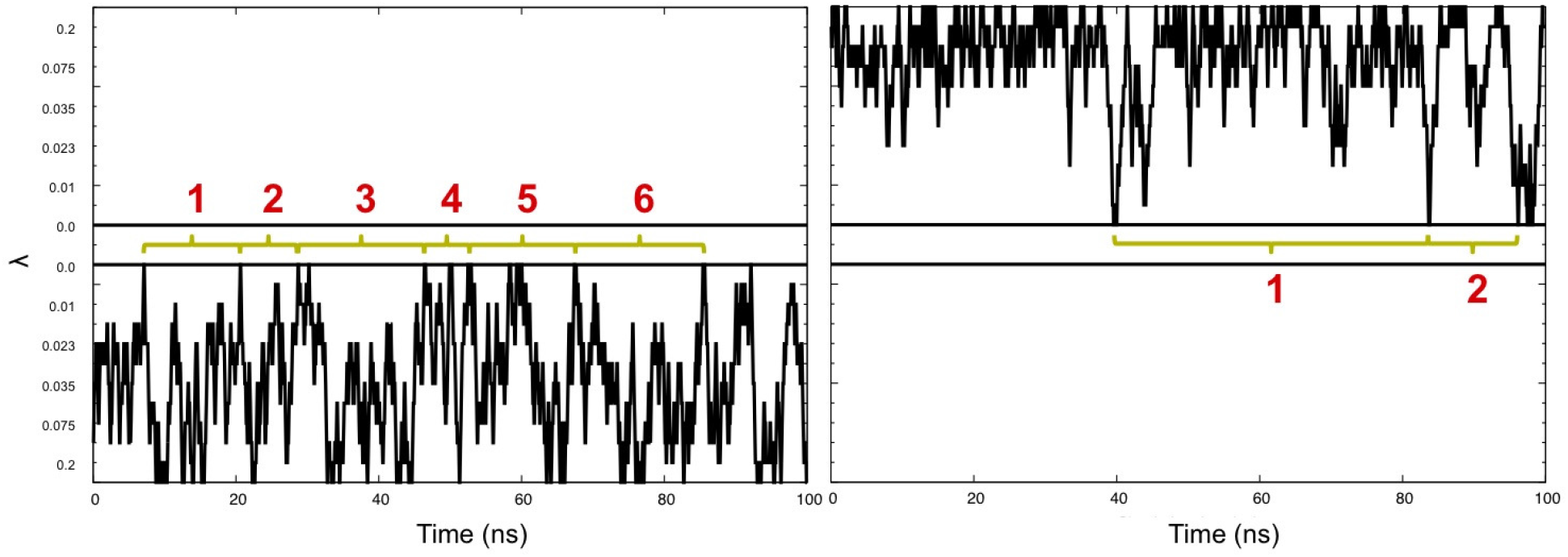
Example of replica walking through λ space, from with a Go-model biasing toward the helix hairpin to a replica with λ = 0 (no bias) (left), and from λ = 0 toward replica with a Go-model biasing to the *β*-barrel (right). Tunneling events are numbered in red. Horizontal black lines mark the λ = 0 replica.

Note that the observed tunneling events cannot be interpreted as folding events leading to either the helix-hairpin or the *β*-barrel state as our RET simulations rely on an artificial dynamics. This is a common problem in all generalized-ensemble and replica-exchange simulations, but one that can be circumvented by reconstructing the free energy landscape of the system under consideration. We show in Figure 3 this landscape projected on the root-mean-square-deviation to either the helix hairpin structure (x-axis) or the *β*-barrel (y-axis). Bin sizes were chosen as 0.8 angstroms, a value smaller than the maximal root-mean-square deviation between models of the 2LCL NMR ensemble, and the landscape was smoothen to interpolate between bins. Note, that this landscape is derived only from the unbiased replica, i.e. the one that has no contribution from a Go-term but is “fed” by the two sides of the ladder of replica, on one side is the physical model biased by the Go-term with varying degrees toward the helix-hairpin, and on the other side is the bias is toward the *β*-barrel.

**Figure 3:**
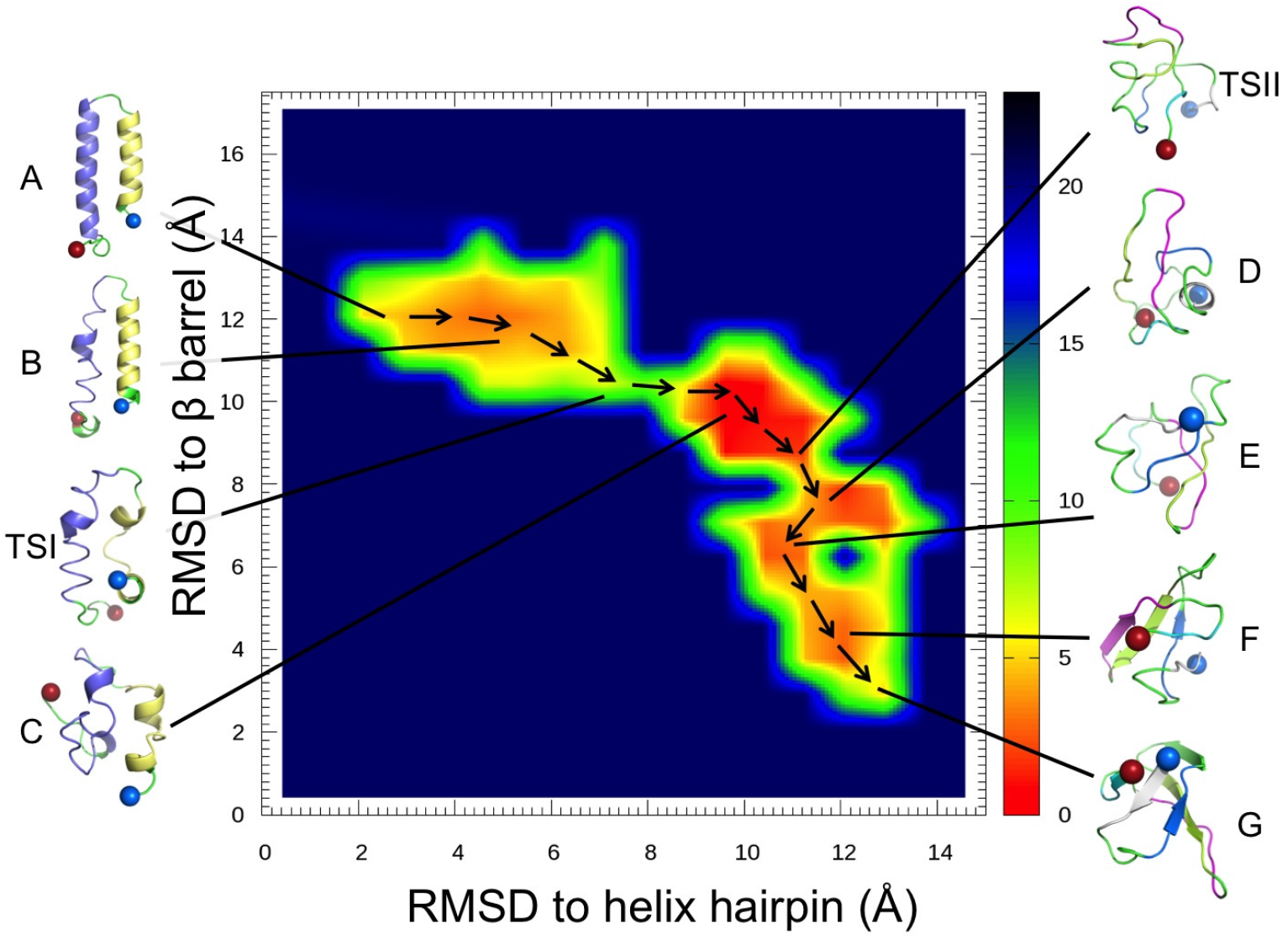
Free energy landscape, in units of RT, of the switching protein RfaH-CTD projected on the root-mean-square deviation (RMSD) with respect to the helix hairpin structure of the protein and with respect to the protein in the *β*-barrel form. Representative configurations are shown for the main basins in the landscape.

Besides the landscape we show also in figure 3 representative structures for the various regions, labeled A to G. Visual inspection and clustering analysis^38^ of the landscape indicates that the *β*-barrel state(region F and G) is the preferred fold of RfaH-CTD, with about 21% of all configurations in the *β*-barrel form. However, the bound state state (region A and B) is also significantly populated, with roughly 6% of configurations in the helix hairpin state. Both folds differ by only approximately 2 RT in free energy, but are separated by a barrier of at least 10 RT. The majority of sampled configurations, 73 %, are either disordered or not representative of either fold. Visual inspection of the free energy landscape in Figure 3 suggests that the transition between helix-hairpin and *β*-barrel state might involve a disordered transition state, see the chain of arrows in the landscape used to trace a possible transition pathway.

Moving from the region of fully-formed helix hairpin (A), helix 2 of RfaH-CTD begins to deteriorate as seen by visual inspection of the configurations in region (B). Moving further along this region, helix 1 begins also to dissolve. Moving out of this basin requires to dissolve the backbone hydrogen bonds that stabilize the two helices, leading to a free energy barrier of about 10 RT, see region TS1. This is supported by the hydrogen-bond analysis in Figure 4 where we define a hydrogen bond by donor acceptor distances of less than 3.5 angstroms and an α angle of less than 30°.

**Figure 4:**
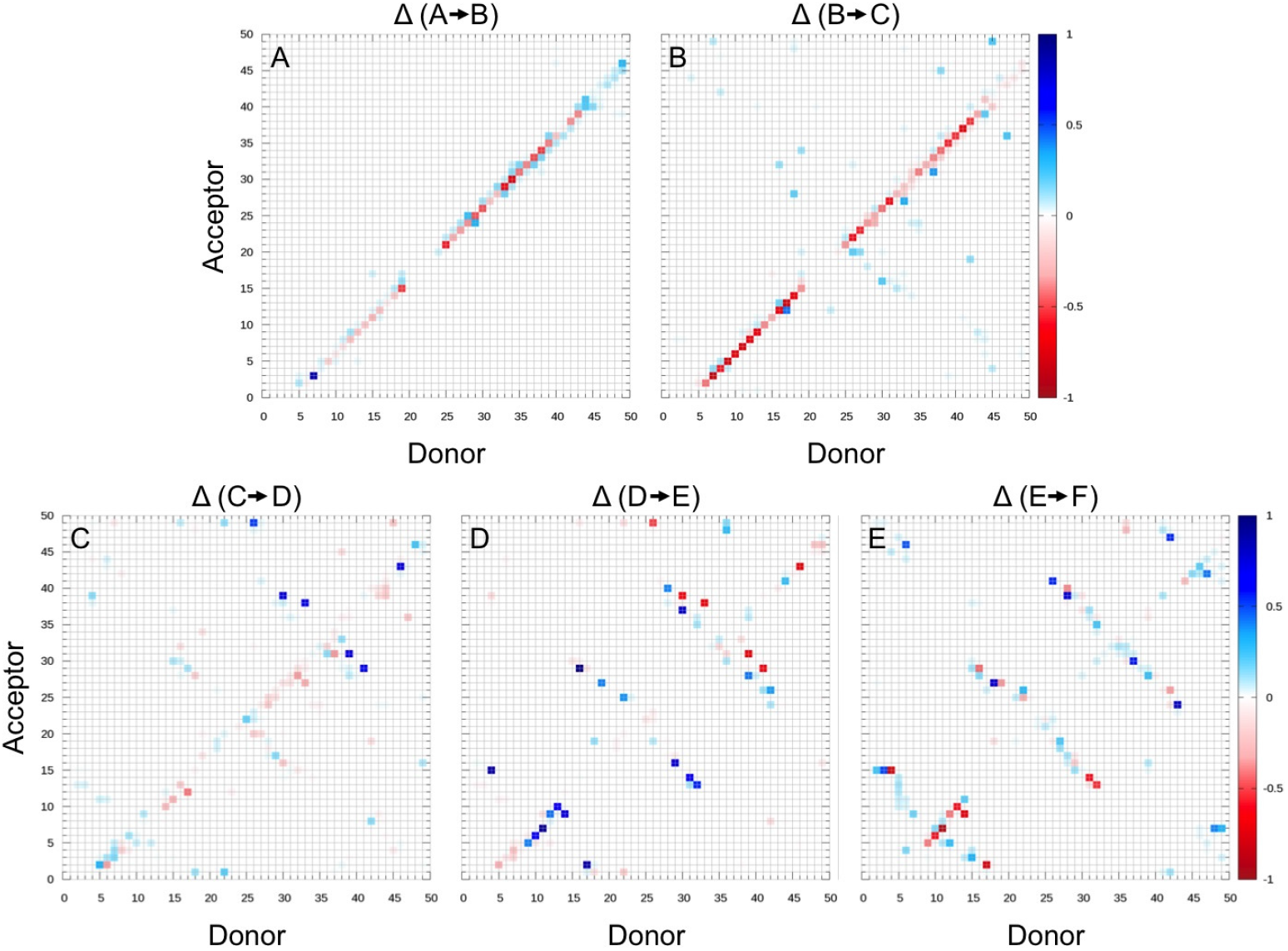
Percentage of structures that gained (blue) or lost (red) backbone hydrogen bonds between the residues indicated on x and y axis, when transitioning from one region of the free energy landscape to another. The involved regions are indicated at the top of each panel, with the indices corresponding to the ones defined in Figure 3.

Upon crossing the barrier (TS1), the RfaH-CTD molecule moves through an ensemble of disordered configurations with little or no defined secondary structure. However in this region (C), *β*-hairpins begin to form in what appears to be a random fashion, and eventually, after crossing a much smaller barrier (TS2) of about 4 RT, a stable hairpin between *β*3 and *β*4 forms in region D. At this point *β*1 has also begun to make contacts with *β*2. Upon entering region E, *β*2 starts to attach to *β*3 of the *β*-hairpin structure, bringing *β*1 with it. Interestingly, there exists a small helix, stabilized by several hydrogen bonds, in the linker region connecting *β*1 and *β*2. This helix positions *β*1 higher up than in the ideal fold likely making it difficult for *β*5 to lay on top of it. Only after finally crossing a third much smaller barrier of less than 2 RT, possibly due to loss of hydrogen bonds in the small helix between *β*1 and *β*2, does RfaH-CTD start to assume in region F the *β*-barrel form. Surprisingly the contacts between *β*5 and *β*4 are maintained in this step where the upper portion of *β*4 bends down slightly allowing *β*5 to lay on top of *β*1 but facing in the wrong direction. In the final step, *β*5 works its way around the N-terminus, thus completing the *β*-barrel (G). This chain of events is again supported by the hydrogen bond analysis of Figure 4. Note that this chain of events is also observed in the tunneling events that we show in Figure 5. While such tunneling events do not necessarily represent “true” transition paths (as they rely on an artificial dynamics), they are added here for illustration.

**Figure 5:**
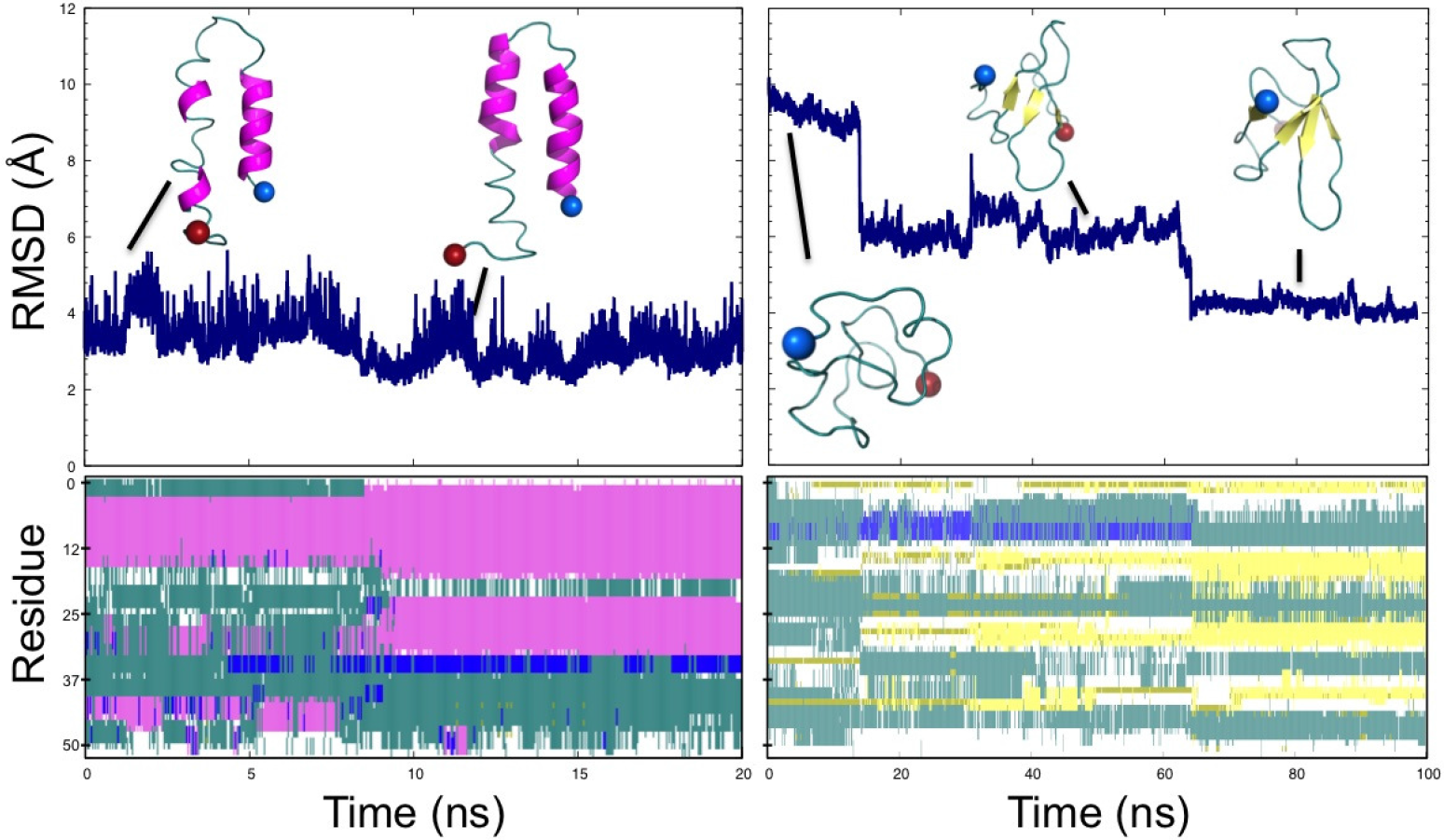
Formation of the helix hairpin (left panels) and the *β*-barrel (right panels). Top panels show the root-mean-square-deviation (RMSD) to the helix hairpin (left) and the *β*-barrel (right). Configurations from various time points are shown. Secondary structure analysis by the VMD program^39^ is shown in the bottom panel. Here, pink represents *α* helices and yellow *β*-sheets.

The above pathway is similar to the one proposed in previous work^11^ that relied on regular REMD simulations. One difference is that in this earlier work, helix 1 breaks first as opposed to our simulations where helix 2 is the one that starts dissolving first. However, in the earlier work, the transition pathway was obtained by following a single replica moving through temperature space. As at high temperatures a replica can cross barriers insurmountable at low temperatures, the observed transition pathways results from an artificial dynamics and does not necessarily describes the correct path. On the other hand, our scenario follows from interpreting the free energy landscape of the protein, not from observed trajectories (which would also result from an artificial dynamics and therefore not necessarily describing the correct pathway). In addition, our scenario is also supported by a comparison of the root-mean-square-fluctuation (RMSF) of residues at the two termini. The C-terminus of RfaH-CTD has a large tail consisting of seven residues following helix 2, while at the N-terminus only three residues precede helix 1. The RMSF values in Figure 6 indicate that the larger tail at the C-terminus is much more flexible than the short end at the N-terminus whose three residues are stabilized by hydrogen bonds with residues 5 and 39 of helix 1 and helix 2. The increased mobility of the seven C-terminal residues adds extra strain on helix 2, thus disrupting its hydrogen bond pattern as is seen in Figure 4, and from visual inspection of clustering data for region (B). Note that this interpretation would not apply to the full-size RfaH protein (instead of only the C-terminal domain RfaH-CTD) which has a much larger linker region preceding helix 1.

**Figure 6:**
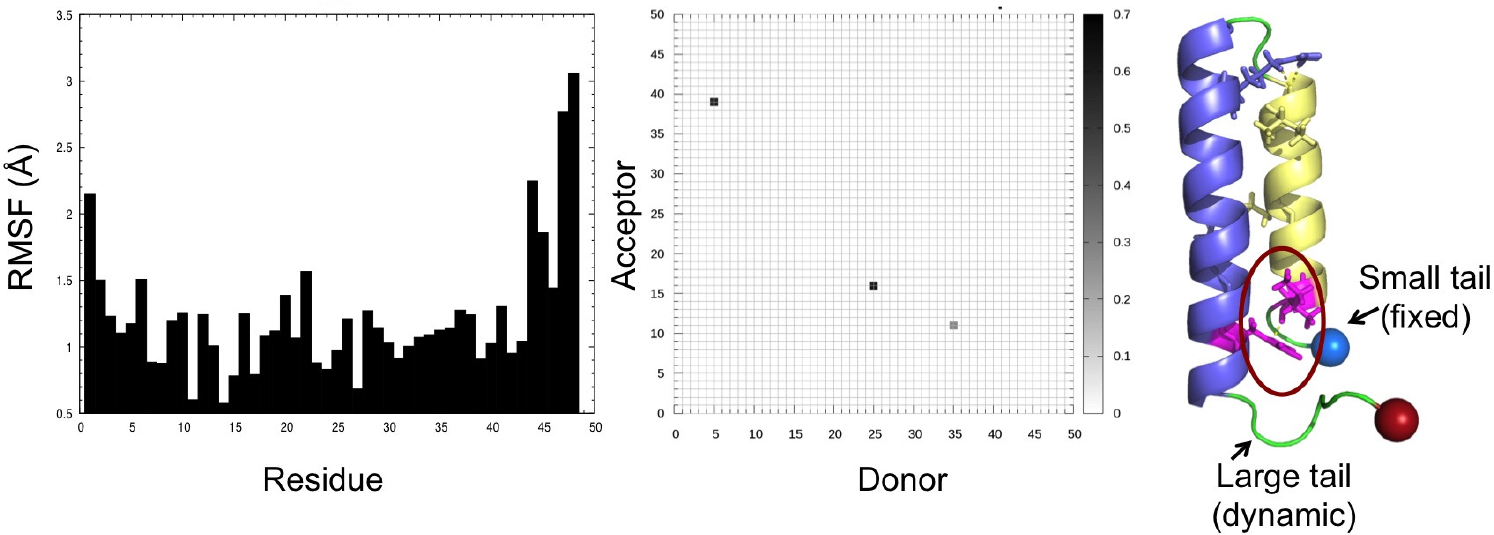
Root-mean-square-fluctuation (RMSF) computed for configurations close to the the helix-hairpin state (less than 2.5 angstroms to the ideal fold) (left panel). Hydrogen bonding between helix 1 and helix 2 for these configurations (middle panel). The helix hairpin structure is shown in the right panel, with the tail segments labeled and the N-terminal tail stabilizing hydrogen bond circled in red.

The differences are much smaller in the remaining parts of the conversion process. Previous work also found that dissolution of the helices is followed by a disordered transition state which precedes formation of a *β*-hairpin between *β*3 and *β*4 and then addition of *β*2. The strand *β*1 was suggested to take longer to align with the developing *β*-sheet due to a larger linker region but once formed would provide a template for addition of *β*5, thus completely folding the barrel. This order of *β*-sheet formation is the same as in our scenario, with the caveat that in our picture the process appears to be more dynamic: *β*-strands continue to grow and re-arrange as additional strands attach to the initial *β*-hairpin, as seen in the panels E-F of figure 4, and become fully-formed only late in the folding pathway toward the *β*-barrel.

## Conclusions

Using a variant of Replica-Exchange-with-Tunneling (RET) we have studied the fold switching process of the 66-residue C-domain RfaH-CTD of the transcription factor RfaH. Our enhanced sampling method allows us to calculate the free energy landscape of the protein projected on suitable coordinates. Analyzing this landscape we propose a mechanism for the conversion process between the helix-hairpin form seen when RfaH-CTD is bound to the he N-terminal domain of RfaH and blocks transcription, and the *β*-barrel form seen in the unbound RfaH-CTD which promotes translation. Consistent with experiments we find that the *β*-barrel is the preferred fold for the isolated RfaH-CTD. However, its free energy is only marginally lower than the helix-hairpin seen in the bound RfAH, but both folds are separated by large barriers resulting from the main chain hydrogen bonds of the helix hairpin. Upon dissolution of the helix hairpin, RfaH-CTD transitions through a disordered state, before a *β*-hairpin forms between *β*3 and *β*4. Later *β*2 attaches to *β*3 of this hairpin, with *β* 1 being in contact already with *β*2. In the final steps, *β*4 bends slightly trapping temporarily *β*5 on top of *β*1 before this strand rearranges and completes the *β*-barrel fold. While the overall pathway is similar to earlier work using traditional REMD simulations^11^, our improved sampling method adds important detail, showing a less structured conversion process with secondary structure only forming late in the process. Together with our earlier work, these results establish the usefulness of our approach for studying switching proteins. We intend now to use our simulation protocol for the simulation of larger switching proteins such as the 93-residue lymphotactin^7^ that would be difficult to study with regular REMD.

## Acknowledgments

The simulations in this work were done using the SCHOONER cluster of the University of Oklahoma and XSEDE resources allocated under grant MCB160005 (National Science Foundation). We acknowledge financial support from the National Science Foundation under grant CHE1266256 and the National Institutes of Health under grant GM120578 and GM120634.

## Supporting Information

A pdf-file containing supplemental figure SF1 (convergence of simulation) and supplemental table ST1 (acceptance rates for exchange moves).

